# Detection of disease-specific signatures in B cell repertoires of lymphomas using machine learning

**DOI:** 10.1101/2023.10.05.561150

**Authors:** Paul Schmidt-Barbo, Gabriel Kalweit, Mehdi Naouar, Lisa Paschold, Edith Willscher, Christoph Schultheiß, Bruno Märkl, Stefan Dirnhofer, Alexandar Tzankov, Mascha Binder, Maria Kalweit

## Abstract

The classification of B cell lymphomas - mainly based on light microscopy evaluation by a pathologist - requires many years of training. Since the B cell receptor (BCR) of the lymphoma clonotype and the microenvironmental immune architecture are important features discriminating different lymphoma subsets, we asked whether BCR repertoire next-generation sequencing (NGS) of lymphoma-infiltrated tissues in conjunction with machine learning algorithms could have diagnostic utility in the subclassification of these cancers. We trained a random forest and a linear classifier via logistic regression based on patterns of clonal distribution, VDJ gene usage and physico-chemical properties of the top-n most frequently represented clonotypes in the BCR repertoires of 620 paradigmatic lymphomas - nodular lymphocyte predominant B cell lymphoma (NLPBL), diffuse large B cell lymphoma (DLBCL) and chronic lymphocytic leukemia (CLL) - as well as 291 control tissues. With regard to DLBCL and CLL, the models demonstrated optimal performance when utilizing only the most prevalent clonotype for classification, while in NLPBL - that has a dominant background of non-malignant bystander cells - a broader array of clonotypes enhanced model accuracy. Surprisingly, the straightforward logistic regression model performed best in this seemingly complex classification problem, suggesting linear separability in our chosen dimensions. It achieved a weighted F1-score of 0.84 on a test cohort including 125 cases from all three lymphoma entities and 58 healthy individuals. Together, we provide proof-of-concept that at least the 3 studied lymphoma entities can be differentiated from each other using BCR repertoire NGS on lymphoma-infiltrated tissues by a trained machine learning model.

**Author Summary:** Lymphoma, a complex group of malignant blood cancers, poses a significant diagnostic challenge due to its diverse subtypes. Yet, precise classification is crucial for tailored treatment. In our research, we developed a machine learning algorithm and conducted comprehensive validation to discern distinct B cell lymphoma subtypes. We therefore leveraged B cell repertoires of lymphoma-infiltrated tissue, as ascertained through next-generation sequencing. Our data offers three key insights: We detail the creation and training of our machine learning algorithm, explaining how we selected features and designed the model. We demonstrate the algorithm’s diagnostic precision using sequencing data from a test-set of patients. Moreover, through a deep dive into the most distinguishing aspects of our algorithm, we unveil distinctive disease-related patterns present within the malignant B cell and its surrounding environment. This analysis showed that both the malignant lymphoma cell, but also healthy bystander immune cells contribute to the distinctive architecture that characterizes a specific lymphoma subtype. We hope our work will contribute towards creating tools to diagnose lymphoma more easily and accurately ultimately leading to better outcomes for patients with this type of cancer.

## Introduction

B cells are one of the essential pillars of the adaptive immune system that generates highly specific and also long-lasting immunity (1, 2). They originate from hematopoietic precursor cells in the bone marrow and acquire their characterizing feature – the B cell receptor (BCR) – in a multistep recombination and selection process (3, 4). Each BCR consists of an unique configuration of paired immunoglobulin heavy (IGH) and light (IGL) chains that mediate antigen binding specificities and upon engagement trigger, in concert with coactivator molecules, a cascade of signaling events that result in activation and proliferation (5, 6). Activated B cells can then differentiate into plasma cells, which produce and secrete immunoglobulins or antibodies to neutralize cognate antigen, or long-lived memory B cells that are capable to quickly mount high-affinity recall responses (7). To guarantee an adequate arsenal of binders for the enormous breadth of foreign antigens, the immune system uses the process of immunoglobulin VDJ recombination to generate maximal sequence diversity (3, 8). During VDJ-recombination in developing immature B cells, randomly chosen variable (V), diversity (D) and joining (J) gene segments within the immunoglobulin loci are recombined to chromosomal sequences encoding a functional BCR (8). This recombination process is facilitated via induced double-strand breaks and DNA repair/ligation mechanisms that may result in additional deletions or insertions that further increase sequence variance of single BCRs (8). On the repertoire level, most of the immunoglobulin diversity is generated in the complementarity-determining region 3 (CDR3) sequence which spans the joined VDJ regions (9, 10). In addition, BCR diversity is boosted by somatic hypermutation (SHM), an iterative affinity maturation process that is initiated in response to antigen in the germinal centers (GCs) of secondary lymphoid tissues (7, 11). These transient but highly specialized microanatomical structures provide a dynamic environment that enables the proper coordination of repeated SHM and selection cycles to evolve polyreactive low-affinity BCRs into antibodies with maximum epitope selectivity (6, 7, 11).

Lymphomas represent hematological neoplasms of differentiated lymphocytes which typically originate in lymphatic tissue (12). Interestingly, 95% of all lymphomas are found in B lineage cells (13). Lymphomas are classified based on histopathological and clinical features that mostly depend on the putative cell of origin that has undergone malignant transformation (14). The BCR takes a prominent role as oncogenic driver and mediator of sustained growth in these malignancies (14). This is not only exemplified by the fact that a specific VDJ-rearrangement characterizes the malignant clonotype in a given patient but also by the acquisition of mutations that mimic chronic active BCR signaling (15). Moreover, comparative analyses of a large number of prior sequencing studies have revealed prominent repertoire restrictions up to the extent of very similar or even identical CDR3 sequences across different patients with the same disease that are now recognized as subtype-specific sequence features for classification (16–18). Furthermore, non-malignant bystander cells occupy varying space within the heterogenous tumor microenvironments of different lymphomas (19, 20). Some lymphomas bear very dominant malignant clonotypes, while in other lymphomas the malignant clonotype is less frequently found in a background of non-malignant bystander cells. These bystander immune cells such as T cells and B cells show a diverse range of other T-cell receptor-gene- or VDJ-rearrangements.

It is generally accepted that the correct diagnostic evaluation of lymphomas by the pathologist is complex in light of the numerous WHO-defined – sometimes rare – entities. Since recognition of the patterns of malignant and bystander cells needs a lot of expertise, evaluation by a consultation pathologist represents a standard in most centers. Here, we hypothesized that high-throughput analysis of the B cell architecture of different types of lymphomas may provide an additional basis for diagnosing disease subtypes. In this context, we set out to employ machine learning algorithms, which have demonstrated their capability to enhance the classification of immune states in antigen receptor-repertoire sequencing data, as valuable tools (21–25). Since BCR next-generation sequencing (NGS) technology relies on unbiased amplification of all VDJ-rearrangements in the tissue of interest, also benign bystander lymphocytes are detected. Here, we present proof-of-concept that a logistic regression model is capable of differentiating between three paradigmatic lymphomas - nodular lymphocyte predominant B cell lymphomas (NLPBL), diffuse large B cell lymphoma (DLBCL), and chronic lymphocytic leukemia (CLL). This data shows the great potential of finding signatures in such large repertoire datasets consisting of the malignant clonotype and benign bystander cells that – clinically – until now remain largely unexploited.

## Results

### Characteristics of the lymphoma and control cohorts

In our study, we analyzed a total of 620 lymphoma cases consisting of 90 NLPBL (formerly nodular lymphocyte predominant Hodgkin lymphoma, NLPHL (26)), 182 DLBCL, and 348 CLL. We enriched the data with 291 control BCR repertoires derived from the blood of healthy donors (HD). Basic characteristics of the cohort are outlined in Table 1.

**Table 1:** Sample characteristics.

| Disease entity |  |  |
| --- | --- | --- |
| HD | Number of samples<br>Type of samples<br>Median age<br>Sex distribution | 291<br>100% PB<br>40 y<br>49% female, 51% male |
| NLPBL | Number of samples<br>Type of samples<br>Median age<br>Sex distribution | 90<br>2% BM, 85% LN, 1% SPL, 12% TM<br>37.5 y<br>30% female, 70% male |
| **DLBCL** | Number of samples<br>Type of samples<br>Median age<br>Sex distribution | 182<br>1.6% LV, 94.6% TM, 3.8% LN<br>71 y<br>46% female, 54% male |
| **CLL** | Number of samples<br>Type of samples<br>Median age<br>Sex distribution | 348<br>100% PB<br>66 y<br>30% female, 70% male |

□ PB = peripheral blood, TM = tumor, LN = lymph node, SPL = spleen, LV = liver, BM = bone marrow

### Broad repertoire metrics in the lymphoma cohorts

In a first step, we compared general repertoire metrics between the four different groups. Blood BCR repertoires of HD were the most diverse and less clonal (Figure 1A-C). Of the lymphoma cases, NLPBL patients showed lowest clonality, followed by patients with DLBCL and CLL (Figure 1A-C). The percentage of somatically hypermutated clonotypes gives a rough estimate of how many clonotypes in the repertoire have undergone antigenic selection. Mean somatic hypermutation in the BCR repertoires was 19.2% in HD, 18.4% in NLPBL, 44.9% in DLBCL and 22.4% in CLL (Figure 1D).

**Figure 1:**
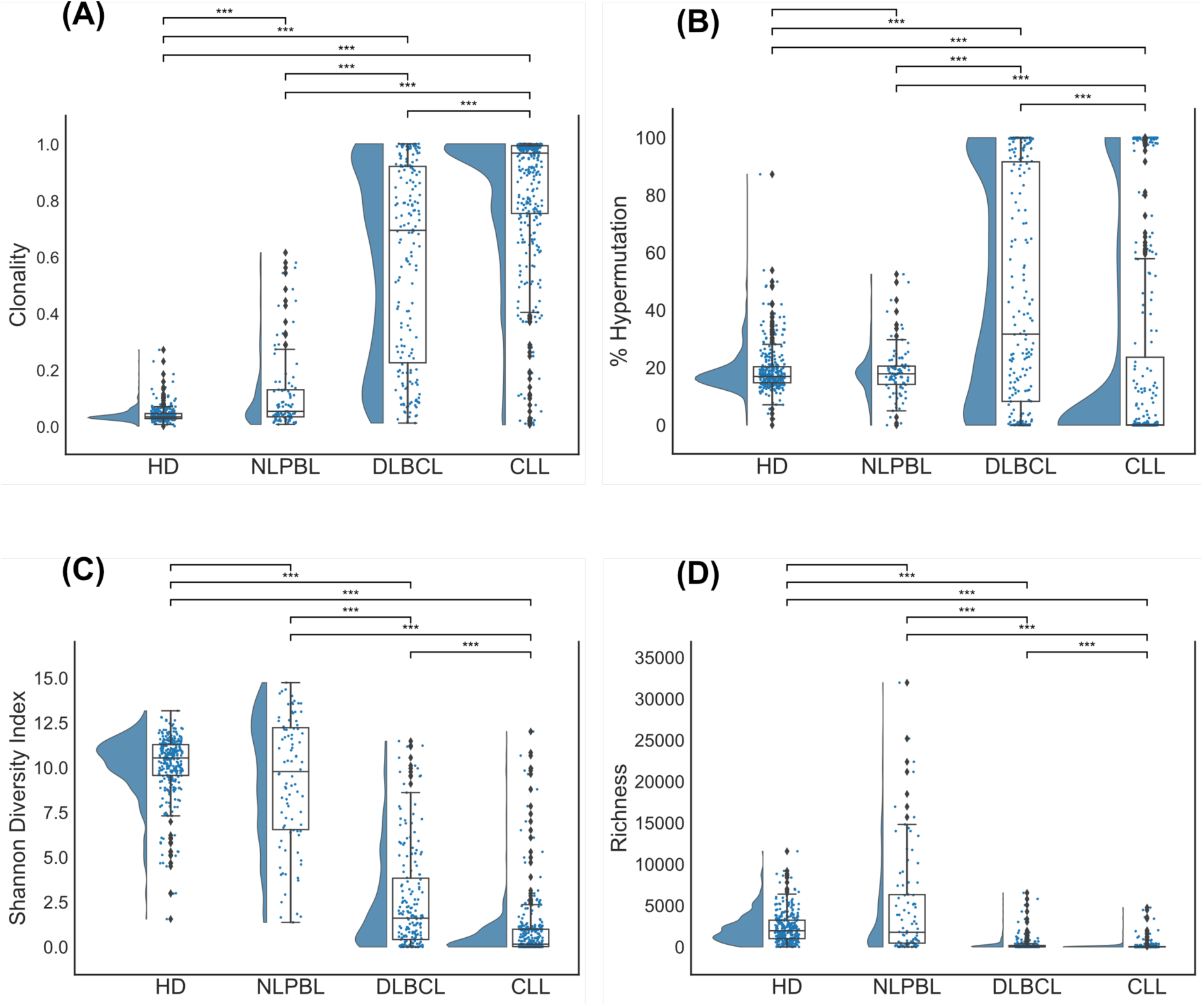
Broad BCR repertoire metrics in lymphoma and control cohorts. In the four panels, the essential repertoire metrics (A) clonality, (B) somatic hypermutation rate, (C) diversity, and (D) richness are shown with corresponding quantiles (Q0.25, Median, Q0.75). We pairwise performed a two-sided Mann-Whitney-U-Test with α = 0.05 (*** p < 0.001).

### Training of machine learning models on BCR repertoires of lymphoma tissue

In our pursuit of accurately classifying lymphoma subtypes based on BCR repertoire characteristics and as an initial benchmarking effort, we developed and trained two machine-learning models. In consideration of interpretability, we opted for two well-established and straightforward models: Logistic regression and random forest. These models were trained using a comprehensive feature set encompassing various aspects of clonotypes, including clonotype fractions, CDR3 sequence lengths, VDJ genes, and Kidera factors (27). Additionally, we incorporated repertoire metrics such as clonality, Shannon diversity index, richness and the fraction of somatic hypermutation within each repertoire into the feature set.

We applied different scenarios in which we challenged the models to discriminate between HD, NLPBL, DLBCL and CLL as well as different combinations thereof. For each scenario we started an individual training phase. For each training phase, the entire dataset comprising the individual classes was partitioned into a training subset (80%) and a test subset (20%). The splitting was performed in a stratified manner, ensuring the proportions of classes were maintained in both subsets, thus guaranteeing an unbiased evaluation of the developed models (Table 2). To account for the imbalance of the classes we used random oversampling, resampling all classes but the majority class.

**Table 2:** Numbers of BCR repertoires used for training.

|  | HD | NLPBL | DLBCL | CLL |
| --- | --- | --- | --- | --- |
| Training | 233 | 72 | 145 | 278 |
| Test | 58 | 18 | 37 | 70 |

Within the training phase, we implemented a robust model optimization strategy. This incorporated a stratified k-fold cross-validation approach with k set to 3 folds. The stratification within the cross-validation approach ensured that the ratio of classes remained consistent across each fold and throughout the entire model training process. A grid search strategy was employed for hyperparameter tuning, traversing a predefined hyperparameter space. A list of resulting hyperparameters and the corresponding search spaces can be found in Supplemental Table 1. To assess the impact of bystander cells, we varied the number of top clonotypes considered in the calculation, ranging from 1 to 10, and subsequently extending to 20, 50 or 100 clonotypes.

### Validation of machine learning algorithm

During the training phase, the models were trained on ⅔ and validated on ⅓ of the training data. This form of validation is crucial in affirming the reliability, predictive capacity, and effectiveness of the models in differentiating the subsets based on the featured BCR repertoire (28). The validation assessment was undertaken through the analysis of key metrics including accuracy, recall, precision and the F1 score. The evaluation of these performance metrics provided a comprehensive understanding of each model’s ability to correctly classify the different cases.

Figure 2A illustrates the averaged validation results achieved by the best performing models for different numbers of clonotypes included in the calculation. While both models, the random forest and the logistic regression, appeared to achieve accurate classification of CLL, DLBCL, and HD cases on the validation sets using information from the top two to three clonotypes alone, logistic regression exhibited improved performance with an increasing number of top repertoire clonotypes for all other cases. However, its performance began to decline when utilizing 50 or 100 clonotypes. In all comparisons including NLPBL cases, logistic regression demonstrated superior performance to random forest when considering information from the top 4-20 clonotypes. The observation that especially discriminating NLPBL benefits from the inclusion of a larger number of top repertoire clonotypes aligns well with the recognized significance of the bystander lymphocyte repertoire in this entity.

**Figure 2:**
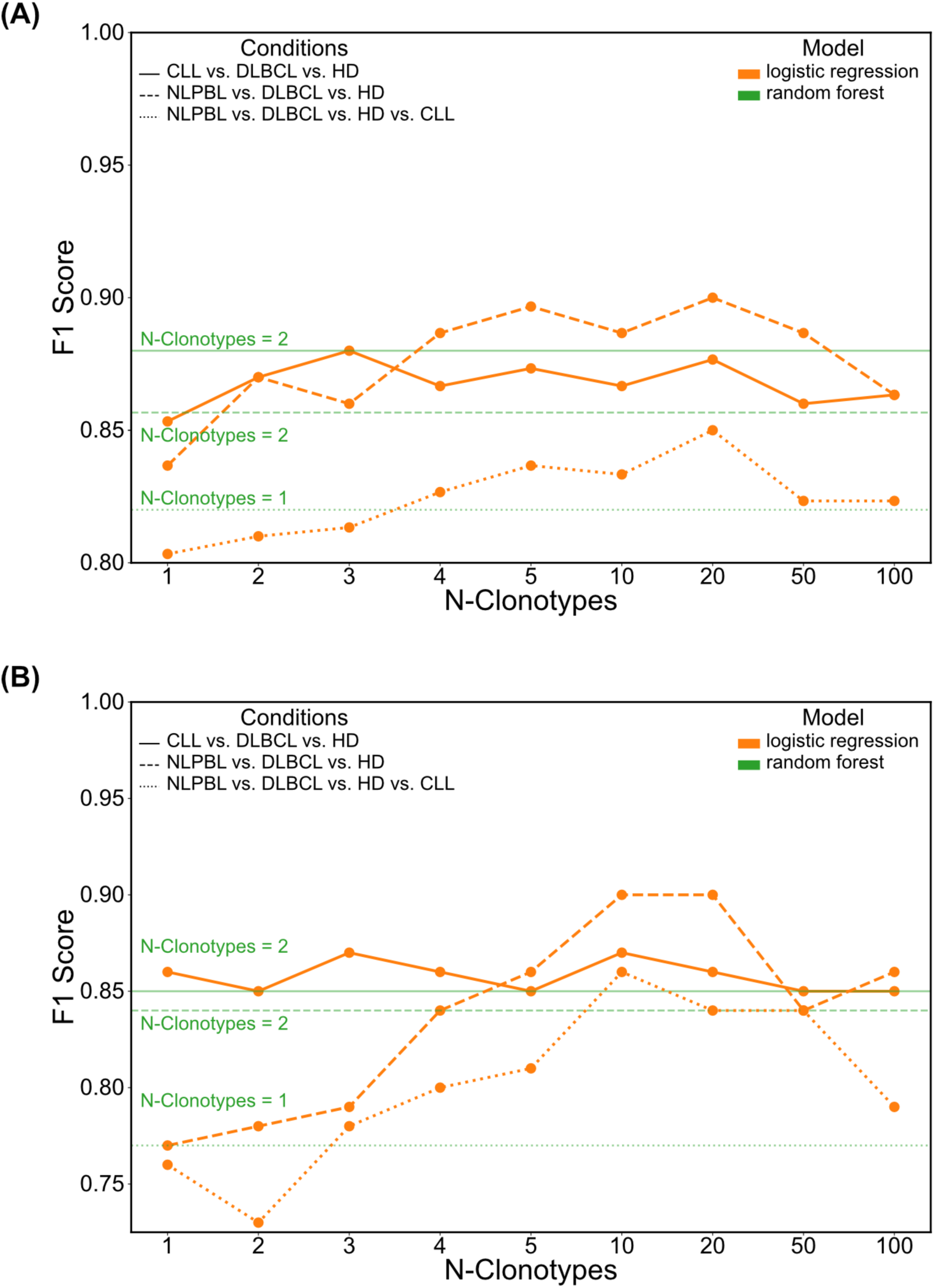
F1 scores of logistic regression models in training and test sets. (A) shows F1 scores averaged over validation folds during training in all three scenarios and (B) those of best validated logistic regression model on the test set in all three scenarios. As a comparison, the performance of the best random forest is displayed in green.

### Final testing of the machine learning models

Upon completing the training phase, the model that exhibited the highest performance based on validation results was further trained on the entire training dataset and evaluated on a previously unseen test dataset. Table 3 showcases crucial metrics, encompassing accuracy, recall, precision, and the F1 score.

**Table 3:** Validation of best models on the independent test set.

| **HD vs. DLBCL vs. CLL**<br>Logistic Regression n-clonotypes = 3 | Precision | Recall | F1 | N |
| --- | --- | --- | --- | --- |
| CLL | 0.82 | 0.94 | 0.88 | 70 |
| DLBCL | 0.85 | 0.59 | 0.70 | 37 |
| HD | 0.95 | 0.97 | 0.96 | 58 |
| Accuracy | 0.87 |  |  |  |
| Weighted Avg. F1 | 0.87 |  |  |  |
| **HD vs. NLPBL vs. DLBCL**<br>Logistic Regression n-clonotypes = 20 | Precision | Recall | F1 | N |
| NLPBL | 0.80 | 0.67 | 0.73 | 18 |
| DLBCL | 0.94 | 0.86 | 0.90 | 37 |
| HD | 0.91 | 1.00 | 0.95 | 58 |
| Accuracy | 0.90 |  |  |  |
| Weighted Avg. F1 | 0.90 |  |  |  |
| **HD vs. NLPBL vs. DLBCL vs. CLL**<br>Logistic Regression n-clonotypes = 20 | Precision | Recall | F1 | N |
| NLPBL | 0.80 | 0.67 | 0.73 | 18 |
| DLBCL | 0.84 | 0.57 | 0.68 | 37 |
| HD | 0.88 | 1.00 | 0.94 | 58 |
| CLL | 0.84 | 0.93 | 0.88 | 35 |
| Accuracy | 0.85 |  |  |  |
| Weighted Avg. F1 | 0.84 |  |  |  |

Figure 2B shows the results of the best validated models on the test data not available during the training phase. Similar to the validation results, we saw superior performance of the logistic regression models. While the first scenario of CLL, DLBCL and HD could be accurately classified by a logistic model only using information of the top three clonotypes, in the other scenarios, the logistic regression heavily benefits from the information of the top 4-20 clonotypes. Although the best performing models with respect to the validation results used 20 clonotypes, models using the information of only 10 clonotypes performed equally or even better on the test set.

### Data separation for n=1 to n=100 clonotypes

To gain a more comprehensive insight into the model’s performance, we conducted Principal Component Analysis (PCA) on our feature list while varying the number of top repertoire clonotypes included in the analysis. Given the clinical relevance of distinguishing between multiple lymphoma subtypes, our primary focus was on the scenario that encompassed all four groups. When visualizing the data in the first two dimensions, we observed overlapping clusters of lymphoma subtypes across different numbers of top repertoire clonotypes (Figure 3A-D). Notably, NLPBL cases exhibited significant overlap with HD repertoires, while DLBCL cases appeared to overlap more with CLL cases. Examining the various embeddings, it became evident that the problem was not perfectly linearly separable along the dimensions of the first and second principal components. However, we did observe the formation of distinct clusters within these two dimensions. Corroborating our performance findings, the most significant separations were achieved when considering the top 4-20 repertoire clonotypes.

**Figure 3:**
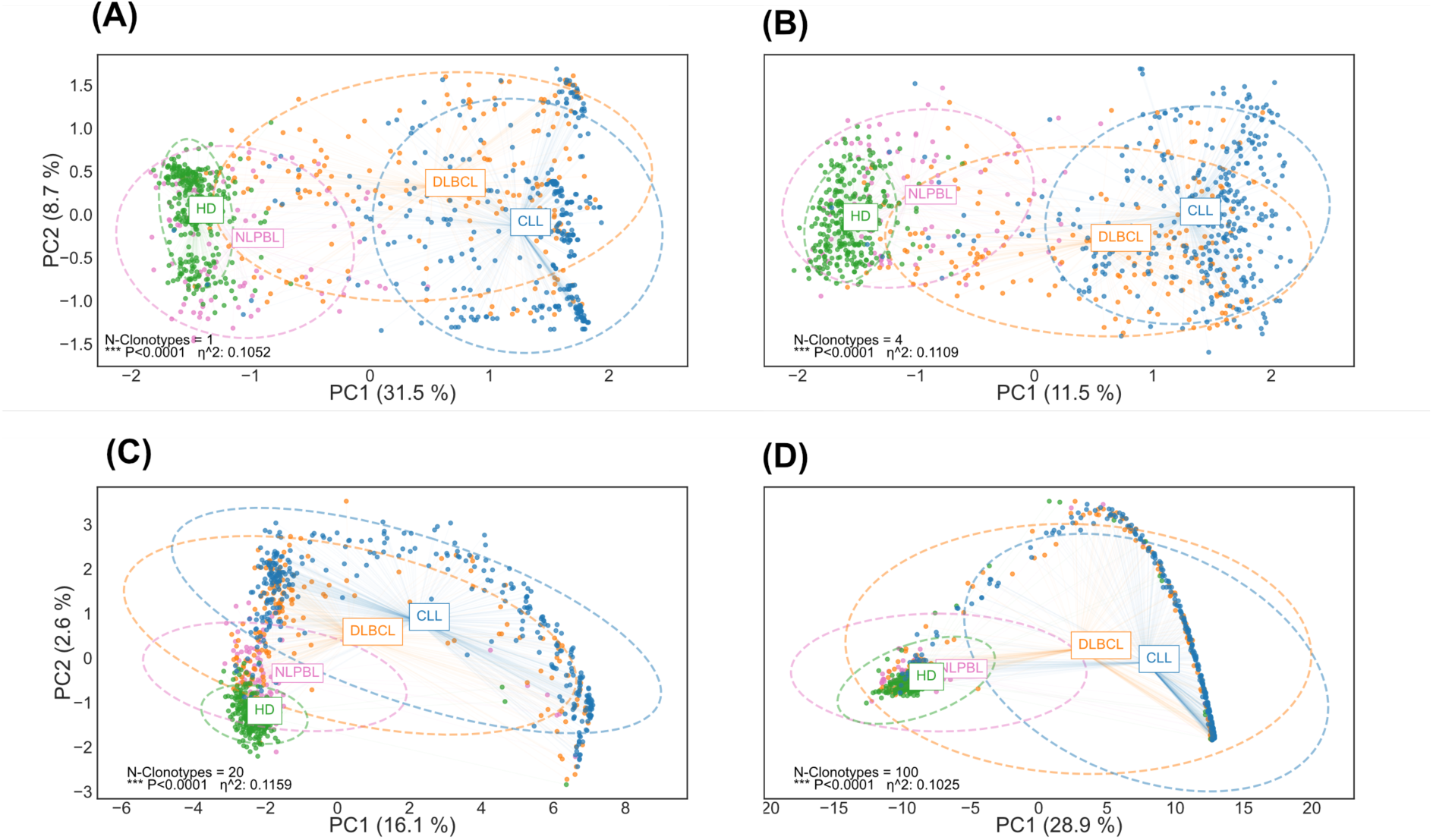
Data separation using n=1 to n=100 top repertoire clonotypes. Principal Component Analysis (PCA) was performed on the feature list while varying the number of top repertoire clonotypes included in the analysis. (A) n=1 clonotype, (B) n=4 clonotypes, (C) n=20 clonotypes, (D) n=100 clonotypes. We compared sample means using a multivariate analysis of variance (MANOVA).

### Exploration of the factors contributing most to correct lymphoma classification

Next, we wished to explore, which of the features contributed most to correct lymphoma classification. We show the 20 predictors with greatest coefficient magnitude averaged over all classes in the best performing model for the comparison of NLPBL, DLBCL with CLL and HD (Figure 4A). This overview helps to understand the overall importance and strength of each predictor variable for the model across all classes. We further analyze the contribution of each predictor between pairs of cohorts (Figure 4B).

**Figure 4:**
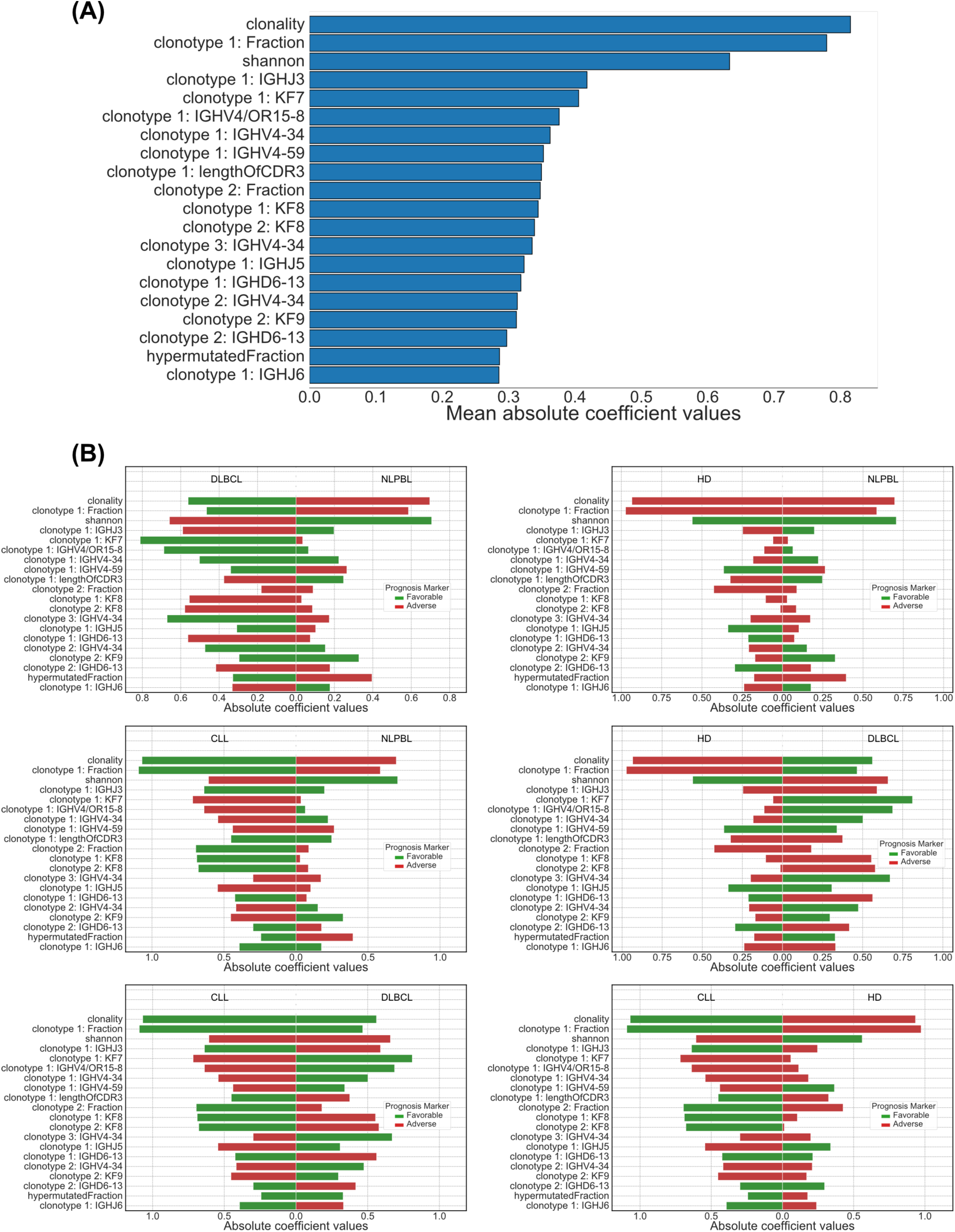
20 lymphoma subset predictors with greatest coefficient magnitude. (A) Predictors were averaged over all classes in the best performing model for discrimination of HD vs. NLPBL vs. DLBCL vs. CLL. (B) Contribution of each predictor to the discrimination between pairs of cohorts.

The three most important predictors of the logistic regression were the frequency of the most abundant clonotype in the repertoire and crude repertoire metrics (clonality, Shannon diversity). While the fraction of somatic hypermutation within a repertoire belonged to the 20 most dominant predictors, repertoire richness seemed to play a minor role in the discrimination between NLPBL, DLBCL, CLL and HD.

Moreover, the model predictions relied on features such as biochemical properties (Kidera factors) and length of the CDR3 sequence of the most dominant clonotypes. Interestingly, we found that Kidera Factor 7, which is associated with flat and extended conformations, and Kidera Factor 9, which represents the partial positive charge of the side chain, served as predictors for DLBCL. In contrast, Kidera factor 8, which reflects the alpha-helical secondary structure, was associated with CLL. The length of the CDR3 of the most dominant clonotype was associated with NLPBL.

Finally, we found specific gene usages of the most dominant clonotypes to add to the discriminative power of the model. For instance, the expression of IGHV4/OR15-8 and IGH4-34 by the most dominant clonotype was predictive of DLBCL and NLPBL, while IGHJ3 was associated with NLPBL and CLL.

## Discussion

We set out to develop a machine-learning tool capable of distinguishing between different types of lymphomas by analyzing the B-cell architecture within lymphoma-infiltrated tissue using BCR-repertoire NGS data. Interestingly, our data revealed that a simple logistic regression with the right set of predictors performed exceptionally well in achieving this goal. It demonstrated a high degree of accuracy when discriminating between the three lymphoma types in our test dataset.

The factors contributing to this successful discrimination were, to some extent, as we anticipated. For example, we found that broad repertoire metrics carried significant discriminatory value. This result aligns with our expectations, especially in diseases characterized by varying levels of immune bystander cells, where such metrics naturally play a role. For this proof-of-concept study, we intentionally selected lymphoma entities characterized by distinct microenvironments. It is conceivable that fitting the algorithm on lymphoma entities with greater similarities could potentially diminish the discriminatory capacity of the basic repertoire metrics, while highlighting the significance of more specific features.

What made our findings particularly intriguing was the discovery that while we saw some discriminative features, which might have been selected based on prior knowledge of the malignant clonotype, some were not as much prioritized by this regression model. For instance, the classical VDJ-rearrangement known to be present in the malignant NLPBL clonotype (V3D3J6) (29, 30) coupled with an unusually long CDR3 sequence length was not prioritized by the algorithm to the extent we would have expected. While one might have anticipated a clear separation based on these features, the discrimination between NLPBL and other lymphomas, such as DLBCL, relied instead quite strongly on other immune repertoire metrics. Overall, most of the discriminatory power came from features of the most dominant clonotypes in the repertoire, yet there was quite some weight on global repertoire metrics underscoring the complexity of lymphoid malignancies. The analysis of patterns of non-malignant bystander cells - unrecognized by traditional bioinformatic analysis - may perspectively shed more light on the immunobiology of lymphoma and may therefore even have broader implications beyond diagnostics.

Several technical aspects of our work need to be discussed. The effective performance of logistic regression and random forest models suggests that the problem at hand may initially appear linearly separable. However, it is noteworthy that logistic regression starts to show limitations when the number of clonotypes exceeds 20, rendering the problem non-linearly separable. This trend prompts the exploration of deep learning models, which are well-suited for handling complex, non-linear relationships within the data. Deep models, such as neural networks, have the capacity to capture intricate patterns and relationships within the data, making them a promising avenue for further exploration. Yet, larger sequencing datasets would be needed to be able to apply deep learning models. In this context, the role of data quantity in optimizing predictive power cannot be overstated. Accumulating more data, particularly diverse and representative samples, is instrumental in bolstering the performance of machine learning models. Additionally, careful consideration of sampling strategies is crucial. Properly balanced and stratified sampling can mitigate issues related to class imbalances and ensure that the model is trained on a comprehensive spectrum of cases. By addressing these aspects, we could refine our machine-learning approach to achieve higher diagnostic accuracy in lymphoma subtyping and potentially even uncover biologically valuable insights into lymphoma clonotypes and their microenvironment.

Our dataset should be considered within the broader framework of evolving research on the application of machine-learning algorithms for lymphoma diagnosis and subtyping. Numerous studies are emerging in the field of digital hematopathology (31–35), and an increasing body of data underscores the pivotal role of machine-learning in harmonizing sequencing data related to lymphoma driver-genes (21–25). Our approach introduces a novel dimension by incorporating immune architecture as an additional layer of analysis. In this respect, the potential of AI may be particularly significant when dealing with challenging scenarios such as suboptimal samples characterized by their small or squeezed nature. In such cases, the BCR repertoire analysis may prove to be less susceptible to interpretation errors compared to traditional morphological assessments. Moreover, specific situations may stand to gain significant benefits, e.g. Richter’s transformation that can sometimes be quite complex to diagnose, while in other instances discerning CLL from DLBCL poses fewer challenges. As an added diagnostic tool in complex cases, it also holds the potential to address the widely recognized shortage of personnel in the field of pathology, which has garnered significant attention within the medical community (36).

Together, our study represents a significant advancement in the field as we demonstrate a compelling proof-of-concept: the ability to differentiate between distinct lymphoma entities by leveraging BCR repertoire NGS on lymphoma-infiltrated tissues through the application of a trained machine learning model. This paves the way for further research and potential clinical applications. In the future, our approach may become a component of digital lymphoma diagnostics as an efficient resource to complement conventional techniques.

## Methods

### Study design

Most of the BCR repertoires have already been deposited in the context of different projects (29, 37–49). The so far unpublished BCR repertoires are deposited together with the previously published ones along with this manuscript to ensure replicability of our results for the comprehensive cohort in the European Nucleotide Archive (ENA) at EMBL-EBI under accession number PRJEB66357 (https://www.ebi.ac.uk/ena/browser/view/PRJEB66357).

New CLL BCR repertoire data has been acquired in a study reviewed by the Ethics Committee of the Martin-Luther-University Halle-Wittenberg after informed written consent (project number 2014-75). The BCR repertoire data of DLBCL samples derives from anonymized paraffin-embedded tissue sections. All studies were conducted in accordance with the ethical principles stated by the Declaration of Helsinki.

### Sample collection, DNA preparation and NGS of BCR repertoires

Peripheral mononuclear cells (PBMC) were isolated from blood of CLL patients or HD by standard density-gradient centrifugation using Ficoll. Genomic DNA was extracted from PBMCs using the GenElute Mammalian Genomic DNA Miniprep Kit (Sigma-Aldrich, St. Louis, USA). In DLBCL and NLPBL patients, DNA was extracted from paraffin-embedded lymphoma-infiltrated tissue as previously described (29).

We used a multiplex PCR based on BIOMED-FR1 primer pool to amplify VDJ rearranged immunoglobulin heavy chain (IGH) loci from 250 ng of genomic DNA. Purified amplicons were pooled at 4 nM, quality-assessed on a 2100 Bioanalyzer (Agilent Technologies) and sequenced on an Illumina MiSeq (paired-end, 2 x 301-cycles, v3 chemistry). Sequence reads were mapped to genomic V, D, J reference sequences using the MiXCR framework(^50^). As reference for sequence alignment, the IMGT library v3 was used. For analysis, we defined each unique complementarity-determining region 3 (CDR3) nucleotide sequence as a clonotype. Non-productive reads and sequences with less than 2 read counts were discarded. IGHV genes that showed ≤ 98% identity to the germline sequence were considered somatically hypermutated. All analyses and data plotting were performed using RStudio (version 1.1.456).

### Immune repertoire metrics

We determined the clonality of the sequenced IGH repertoires using the formula “1-Pielou’s evenness” (51, 52). In our context, evenness quantifies the relative prevalence of distinct B cell types within each repertoire. It is calculated according to the formula J = H’/log2(S) with H’ being the Shannon diversity index (53) and S the total clonotype number (richness) (54) in a distinct sample. A clonality index of 1 indicates that the analyzed sample comprises a single clonotype, while 0 indicates complete clonal diversity.

### Machine Learning

#### Data Preprocessing

For each clonotype in all repertoires, we extracted the clonotype fraction, lengths of CDR3 sequences and the VDJ arrangement. Since VDJ genes are categorical features, we applied one-hot-encoding. Representing the theoretical physico-chemical properties, we calculated the ten Kidera Factors from the individual amino acid sequence of each CDR3 region and augmented the dataset with the clonality, Shannon diversity index and richness for each repertoire as described. In order to deal with the variable number of clonotypes we set a fixed number of clonotypes per repertoire ranging from n=1 to 10 and subsequently extending to 20, 50 or 100 clonotypes. Based on their clonotype fractions, we extracted the n most dominant clonotypes within each repertoire while discarding the remaining clonotypes. For repertoires with fewer than n clonotypes we applied zero-padding to ensure consistent input dimensions across all samples. We concatenated the features of the n clonotypes to a single vector representing a single repertoire within 111 (n = 1) up to 10151 (n = 100 dimensions. Depending on a freely varying hyperparameter we scaled the numerical feature across all vectors to have zero mean and unit variance. We partitioned the data into a training (80%) and test set (20%) ensuring the proportions of classes were maintained in both subsets.

#### Training

Within the training process we fitted two different model types in a supervised manner. The first model was a random forest consisting of an ensemble of single decision trees. The second model consists of a logistic regression minimizing a multinomial loss. Using a stratified k-fold cross-validation approach with k set to 3 folds we fitted each model with a different set of hyperparameters (supplemental table 1) to a subset of the training data. We calculated the performance in form of the F1 score on the remaining part of the training set and averaged the obtained scores over all folds. The best performing settings of hyperparameters were chosen and used to train the model on the entire training set.

#### Testing

The best performing models with respect to the training phase were used to predict unseen data from the separate test set. We compared the predictions of each model to the known labels and calculated a final F1 score.

The whole analysis was implemented in Python 3.11.5 with scikit-learn 1.3.0 using a Conda environment. All calculations were performed on a MacBook Pro using a M1 processor, Kernel Version Darwin 22.6.0 and macOS 13.5.2.

All code is available under https://github.com/paulovic96/immusign.

## Author contributions

Idea & design of research project: MB, MK. Supply of critical material (e.g. patient material, cohorts): BM, MB, LP, CS, EW; Establishment of Methods: MK, GK, MN; Experimental work: PSB, MK; Data analysis and interpretation: MB, MK, GK, PSB, AT, SD; Drafting of manuscript: MB, MK, PSB.

## Acknowledgments

We thank the lymphoma patients and healthy donors for their great support. Moreover, we thank Christoph Wosiek, Aline Patzschke, Jenny Wehde, Yiqing Du and Katrin Nerger for excellent technical assistance. We also thank Roland Mertelsmann for his valuable advice and the CRIION for core AI support. This project was funded by the Mertelsmann Foundation (grant to MB).

---

**Supplemental Table 1:** Hyperparameters used during training

|  | Feature | Values |
| --- | --- | --- |
| General | number of clonotypes | 1, 2, 3, 4, 5, 10, 20, 50, 100 |
|  | standardize data | Yes, No |
| Random forest | number of decision trees | 200, 800 |
|  | maximum depth of trees | 8, 16 |
| Logistic Regression | regularization | L2, L1 |
|  | C | [0.01, 0.1, 1, 10] |

**Final Hyperparameters**
| **HD vs. DLBCL vs. CLL** | Feature | Values |
| --- | --- | --- |
| General | number of clonotypes | 3 |
|  | standardize data | Yes |
| Logistic Regression | regularization | L2 |
|  | C | 10 |
| **HD vs. NLPBL vs. DLBCL** | Feature | Values |
| General | number of clonotypes | 20 |
|  | standardize data | Yes |
| Logistic Regression | regularization | L2 |
|  | C | 0.1 |
| **HD vs. NLPBL vs. DLBCL vs. CLL** | Feature | Values |
| General | number of clonotypes | 20 |
|  | standardize data | Yes |
| Logistic Regression | regularization | L2 |
|  | C | 1 |

## Notes

### Competing Interest Statement

The authors have declared no competing interest.

